# Ronapreve (REGN-CoV; casirivimab and imdevimab) reduces the viral burden and alters the pulmonary response to the SARS-CoV-2 Delta variant (B.1.617.2) in K18-hACE2 mice using an experimental design reflective of a treatment use case

**DOI:** 10.1101/2022.01.23.477397

**Authors:** Lee Tatham, Anja Kipar, Joanne Sharp, Edyta Kijak, Joanne Herriott, Megan Neary, Helen Box, Eduardo Gallardo Toledo, Anthony Valentijn, Helen Cox, Henry Pertinez, Paul Curley, Usman Arshad, Rajith KR Rajoli, Steve Rannard, James Stewart, Andrew Owen

**Author notes:** Both authors contributed equally to the work (Lee Tatham, Anja Kipar).

## Abstract

**Background:** Ronapreve demonstrated clinical application in post-exposure prophylaxis, mild/moderate disease and in the treatment of seronegative patients with severe COVID19 prior to the emergence of the Omicron variant in late 2021. Numerous reports have described loss of *in vitro* neutralisation activity of Ronapreve and other monoclonal antibodies for BA.1 Omicron and subsequent sub-lineages of the Omicron variant. With some exceptions, global policy makers have recommended against the use of existing monoclonal antibodies in COVID19. Gaps in knowledge regarding the mechanism of action of monoclonal antibodies are noted, and further preclinical study will help understand positioning of new monoclonal antibodies under development.

**Objectives:** The purpose of this study was to investigate the impact of Ronapreve on compartmental viral replication as a paradigm for a monoclonal antibody combination. The study also sought to confirm absence of *in vivo* activity against BA.1 Omicron (B.1.1.529) relative to the Delta (B.1.617.2) variant.

**Methods:** Virological efficacy of Ronapreve was assessed in K18-hACE2 mice inoculated with either the SARS-CoV-2 Delta or Omicron variants. Viral replication in tissues was quantified using qRT-PCR to measure sub-genomic viral RNA to the E gene (sgE) as a proxy. A histological examination in combination with staining for viral antigen served to determine viral spread and associated damage.

**Results:** Ronapreve reduced sub-genomic viral RNA levels in lung and nasal turbinate, 4 and 6 days post infection, for the Delta variant but not the Omicron variant of SARS-CoV-2 at doses 2-fold higher than those shown to be active against previous variants of the virus. It also appeared to block brain infection which is seen with high frequency in K18-hACE2 mice after Delta variant infection. At day 6, the inflammatory response to lung infection with the Delta variant was altered to a mild multifocal granulomatous inflammation in which the virus appeared to be confined. A similar tendency was also observed in Omicron infected, Ronapreve-treated animals.

**Conclusions:** The current study provides evidence of an altered tissue response to the SARS-CoV-2 after treatment with a monoclonal antibody combination that retains neutralization activity. These data also demonstrate that experimental designs that reflect the treatment use case are achievable in animal models for monoclonal antibodies deployed against susceptible variants. Extreme caution should be taken when interpreting prophylactic experimental designs when assessing plausibility of monoclonal antibodies for treatment use cases.

## Introduction

A concerted global effort since the emergence of SARS-CoV-2 in late 2019 resulted in a toolbox of putative interventions that were brought through development at unprecedented speed. The rapid development and implementation of vaccination programmes has had a transformational impact on control of the pandemic in some countries but ongoing efforts for vaccine equity continue to be critical.^1^ In addition, first generation antiviral drugs have emerged from repurposed small molecules from other antiviral development programmes, such as drugs originally developed for ebola, influenza or prior coronaviruses. More potent antivirals continue to emerge, but considerable research is still required to optimise deployment of existing agents (including evaluation of regimens composed of drug combinations).^2, 3^

Neutralising monoclonal antibodies targeting the spike protein on the surface of SARS-CoV-2 were also brought forward with commendable speed, but the urgency of the pandemic necessitated that key knowledge was not collected during the accelerated development process. Ronapreve (REGN-COV2) is composed of two such monoclonal antibodies (casirivumab and imdevimab), and demonstrated clinical efficacy against pre-Omicron SARS-CoV-2 variants in post-exposure prophylaxis,^4^ early treatment,^5^ and in the treatment of seronegative patients with severe COVID19.^6^ With successive emergence of Omicron sub-lineages, all approved monoclonal antibodies have lost varying degrees of neutralisation capability such that continued efficacy in all use cases is no longer plausible based upon the current understanding of the pharmacokinetic-pharmacodynamic relationship.^7–9^ As a result, no monoclonal antibodies are currently recommended by the NIH or WHO.^10–12^ Each antibody in Ronapreve exhibits molar potency against previous SARS-CoV-2 variants which are orders of magnitude higher than current repurposed small molecule drugs such as molnupiravir and nirmatrelvir^10, 13^ but they were given in combination to reduce the risk of emergence of resistance as has been widely documented for monoclonal antibodies used against susceptible variants as monotherapy.^14–20^

Several variants of concern (VOC) have emerged over the past 2 years to which at least one of the antibodies in Ronapreve has retained activity *in vitro.^21^* Moreover, studies in K18-hACE2 transgenic mice clearly demonstrated virological efficacy of Ronapreve against previous variants (not including Delta which was not studied).^22^ Several studies have also investigated the activity of casirivumab and imdevimab (alone or in combination) against pseudovirus engineered to express the BA.1 Omicron spike protein or authentic virus.^23–25^ All studies have demonstrated compromised activity of the Ronapreve combination in these assays. However, other studies reported residual activity of the individual antibodies when studied in isolation, albeit with substantially lower activity. Unlike other monoclonal antibodies, extremely high doses of casirivumab and imdevimab (up to 8000 mg intravenously) have been studied safely and pharmacokinetics at these doses far exceed stringent target concentrations developed by the manufacturers for ancestral SARS-CoV-2.^5^

Recent studies provided evidence that intraperitoneal administration of neutralising human antibodies protect K18-hACE2 mice from lung infection and clinical disease in both prophylactic (hours to 3 days prior to intranasal infection, using an ancestral SARS-CoV-2 and the Delta variant) and therapeutic settings (up to 4 days post intranasal infection).^26, 27^ A study using neutralising murine monoclonal antibodies demonstrated significant reduction of viral titres in the lungs at 2 days post infection (dpi) i.e. the peak of lung infection in untreated mice, when mice were treated with the antibody at 6 hours post intranasal infection. Similarly, prophylactic treatment (day −1) prior to infection with an original virus isolate significantly reduced weight loss and viral titres in nasal turbinate, lungs and brain at 5 dpi; interestingly, treatment at 5.5 hours post infection had the same effect on body weight and viral loads in all tested organs except the lungs.^28^

The purpose of this study was to investigate the ability of monoclonal antibody combinations to mitigate pulmonary and neurological manifestations of SARS-CoV-2 infection using Ronapreve and the Delta variant as a paradigm for activity against a susceptible variant. *In vivo* validation of prior *in vitro* assay readouts for neutralisation of BA.1 Omicron by Ronapreve is also presented.

## Methods

### Materials

Materials were purchased and used as received without further purification: chloroform, isopropanol, ethanol, phosphate buffered saline (PBS) and nuclease-free water were purchased from Fisher Scientific (UK). Male K18-hACE2 mice were purchased from Charles River (France). Ronapreve (casirivimab and imdevimab) was kindly provided by Roche (Switzerland). TRIzol^TM^, GlycoBlue^TM^, Phasemaker^TM^ tubes and TURBO DNA-free^TM^ kit were purchased from Fisher Scientific (UK). GoTaq^®^ Probe 1-Step RT-qPCR System was purchased from Promega (US). SARS-CoV-2 (2019nCoV) CDC qPCR Probe Assay was purchased from IDT (US). Precellys CKmix lysing tubes were purchased from Bertin Instruments (France). For immunohistology, a rabbit anti-SARS-CoV nucleoprotein antibody was purchased from Rocklands, the peroxidase blocking buffer and the Envision+System HRP Rabbit and the diaminobenzidine from Agilent/DAKO. All other chemicals and reagents were purchased from Merck (UK) and used as received, unless stated otherwise.

### Virus isolates

The Delta variant (B.1.617.2) hCoV-19/England/SHEF-10E8F3B/2021 (GISAID accession number EPI_ISL_1731019), was kindly provided by Prof. Wendy Barclay, Imperial College London, London, UK through the Genotype-to-Phenotype National Virology Consortium (G2P-UK). Sequencing confirmed it contained the spike protein mutations T19R, K77R, G142D, Δ156-157/R158G, A222V, L452R, T478K, D614G, P681R, D950N. The Omicron variant (B.1.1.529/BA.1) isolate M21021166 was originally isolated by Prof. Gavin Screaton, University of Oxford^24^, UK and then obtained from Prof. Wendy Barclay, Imperial College London, London, UK through the Genotype-to-Phenotype National Virology Consortium (G2P-UK). Sequencing confirmed it contained the spike protein mutations A67V, Δ69-70, T95I, G142D/Δ143-145, Δ211/L212I, ins214EPE, G339D, S371L, S373P, S375F, K417N, N440K, G446S, S477N, T478K, E484A, Q493R, G496S, Q498R, N501Y, Y505H, T547K, D614G, H655Y, N679K, P681H, N764K, A701V, D796Y, N856K, Q954H, N969K, L981F. The titres of all isolates were confirmed on Vero E6 cells and the sequences of all stocks confirmed.

### Animal studies

All work involving SARS-CoV-2 was performed at containment level 3 by staff equipped with respirator airstream units with filtered air supply. Prior to the start of the study, all risk assessments and standard operating procedures were approved by the University of Liverpool Biohazards Sub-Committee and the UK Health and Safety Executive.

All animal studies were conducted in accordance with UK Home Office Animals Scientific Procedures Act (ASPA, 1986). Additionally, all studies were approved by the local University of Liverpool Animal Welfare and Ethical Review Body and performed under UK Home Office Project License PP4715265. Male mice (20-30 g) carrying the human ACE2 gene under the control of the keratin 18 promoter (K18-hACE2; formally B6.Cg-Tg(K18-ACE2)2Prlmn/J) were housed in individually-ventilated cages with environmental enrichment under SPF barrier conditions and a 12-hour light/dark cycle at 21 °C ± 2 °C. Free access to food and water was provided at all times.

Mice were randomly assigned into groups and acclimatized for 7 days. Mice in each group were anaesthetised under 3% isoflurane and inoculated intranasally with 100 μL of either 10^3^ PFU of SARS-CoV-2 Delta variant (B.1.617.2) or Omicron variant (B.1.1.529) in phosphate buffered saline (PBS). After 24 hours, mice from each group were treated with a single dose (100 μL) of either the saline control or 400 μg Ronapreve, diluted in saline, via intraperitoneal (IP) injection. All animals were weighed and monitored daily throughout the experiment. At 4 and 6 days following infection, groups of mice were sacrificed via a lethal IP injection of pentobarbitone, followed by cardiac puncture and immediate exsanguination from the heart. Animals were immediately dissected and the right lung as well as fragments from the nasal turbinates collected and frozen at −80°C for RNA extraction. The left lung lobe and the head were fixed in 10% buffered formalin for 48 hours and then stored in 70% ethanol until processing for histological and immunohistological examination.

### Quantification of viral RNA

RNA isolation from lung and nasal turbinate samples, RNA quantification, and DNAse treatment has been detailed previously.^25^

The viral RNA derived from the lung and nasal turbinate samples was quantified using a protocol for quantifying the SARS-CoV-2 sub-genomic E gene RNA (sgE)^29^ using the GoTaq^®^ Probe 1-Step RT-qPCR System (Promega).

Quantification of SARS-CoV-2 E SgRNA was completed utilising primers and probes previously described elsewhere^29^ and were used at 400 nM and 200 nM, respectively (IDT), using the GoTaq^®^ Probe 1-Step RT-qPCR System (Promega). Quantification of 18S RNA utilised previously described primers and probe sequences,^30^ and were used at 300 nM and 200 nM, respectively (IDT), using the GoTaq^®^ Probe 1-Step RT-qPCR System (Promega). Methods for the generation of the 18S and sgE RNA standards have been outlined previously.^31^ Both PCR products were serially diluted to produce standard curves in the range of 5 x 10^8^ - 5 copies/reaction via a 10-fold serial dilution. DNAse treated RNA at 20,000 ng/mL or dH_2_O were added to appropriate wells producing final reaction volumes of 20 μL. The prepared plates were run using a qTOWER^3^ Real-Time PCR Detector (Analytik Jena). Thermal cycling conditions have been detailed previously.^25^ The sgE data were normalised to 18S data for subsequent quantitation.

### Statistical analysis

An unpaired, two-tailed, t-test was used to compare the differences in lung and nasal turbinate viral RNA between the control (saline) and Ronapreve treatment groups at days 4 and 6. A P-value of ≤ 0.05 was considered statistically significant. All statistical analysis was completed using GraphPad Prism version 7.

### Histological and immunohistological analyses

The fixed left lung was routinely paraffin wax embedded. Heads were sawn longitudinally in the midline using a diamond saw (Exakt 300; Exakt) and the brain left in the skull. Heads were gently decalcified in RDF (Biosystems) for twice 5 days, at room temperature (RT) and on a shaker, then both halves paraffin wax embedded. Consecutive sections (3-5 μm) were prepared and stained with hematoxylin eosin (HE) for histological examination or subjected to immunohistological staining to detect SARS-CoV-2 antigen (performed in an autostainer; Agilent), using the horseradish peroxidase (HRP) method and rabbit anti-SARS-CoV nucleocapsid protein (Rockland) as previously described^32^. Briefly, sections were deparaffinized and rehydrated through graded alcohol. Antigen retrieval was achieved by 20 min incubation in citrate buffer (pH 6.0) at 98 °C in a pressure cooker. This was followed by incubation with the primary antibody (diluted 1:3,000 in dilution buffer; Dako) overnight at 4 °C, a 10 min incubation at RT with peroxidase blocking buffer (Agilent) and a 30 min incubation at RT with Envision+System HRP Rabbit (Agilent). The reaction was visualized with diaminobenzidin (DAB; Dako) for 10 min at RT. After counterstaining with hematoxylin for 2 s, sections were dehydrated and glass coverslipped. In selected animals (see Supplemental Table S1), lungs were also stained for CD3 (T cell marker), CD45R/B220 (B cell marker) and Iba1 (macrophage marker), as previously described^32^.

## Results

### Body weight

Weight was monitored throughout the study as a marker for health. Figure 1 shows mouse weights relative to baseline (day 0; prior to SARS-CoV-2 inoculation). All animals displayed weight loss at day 2 post infection (9.3-14.3% of body weight); this was less rapid in the Delta variant infected animals, albeit without statistical significance. Most animals regained some weight (2.1-5.2%) by day 3 and most reached pre-infection levels (around 95%) by day 6 with the exception of the control Delta variant infected animals which showed progressive weight loss after day 4, partly reaching the clinical endpoint (up to 20% weight loss) by day 6.

**Figure 1.**
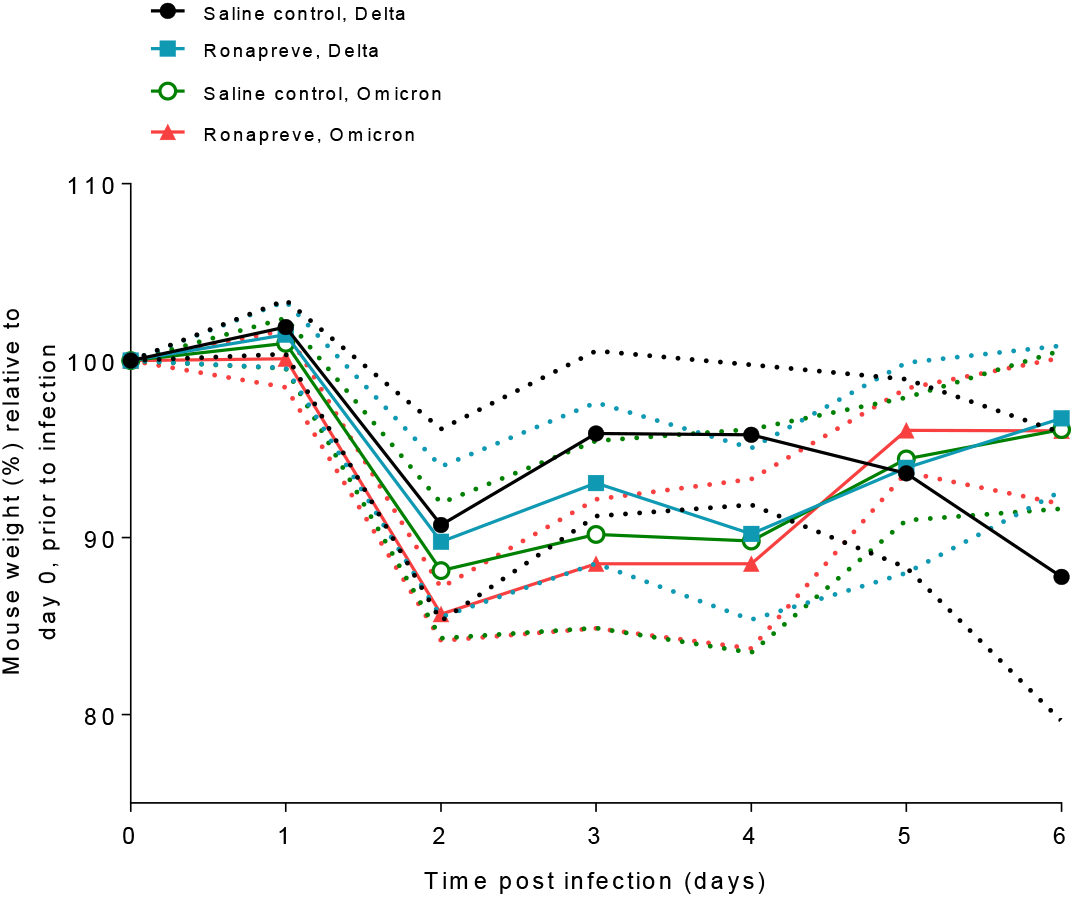
Mouse weights separated by treatment group and infection status. Weights are the percentage of the initial weight recorded at day 0 prior to infection. Standard deviations are indicated by the dashed plots.

### Effect of Ronapreve on viral replication

To determine the viral load in animals infected with each variant and subsequently dosed with either saline (controls) or Ronapreve, total RNA was extracted from the lung and nasal turbinate samples of animals culled on days 4 and 6 post infection. Viral replication was quantified using qRT-PCR to measure sub-genomic viral RNA to the E gene (sgE) as a proxy. The results are illustrated in Figure 2. In SARS-CoV-2 Delta variant infected animals, the amount of sgE RNA was generally reduced after Ronapreve treatment compared to the saline treated mice. At 4 dpi the difference was significant in the nasal turbinates (log10 fold decrease: −0.556, P=0.037) but not in the lung (log10 fold reduction: −0.602, P=0.065), whereas at 6 dpi, the difference was not significant in the nasal turbinates (log10 fold decrease: −1.369, P=0.111) but significant in the lung (log10 fold reduction: −1.667, P=0.033).

**Figure 2.**
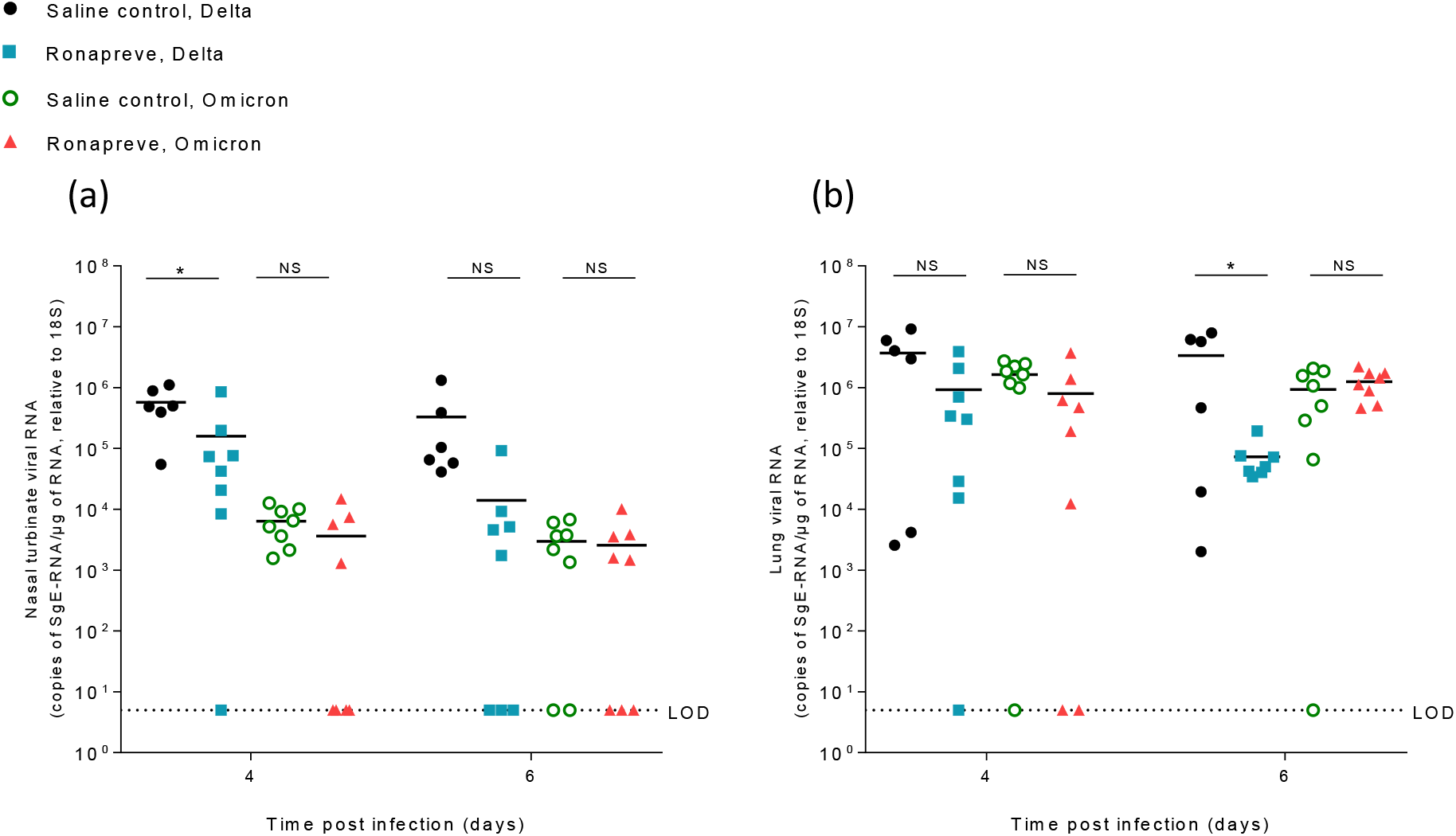
Viral quantification of SARS-CoV-2 sub-genomic RNA (sgE), relative to 18S, using qRT-PCR from nasal turbinate (a) and lung (b) samples harvested from each group on days 4 and 6 post infection. Mice infected with the Delta variant were administered with a single IP dose of either saline (n=12) or Ronapreve, 400 μg/mouse, in saline (n=16). Equally, mice infected with the Omicron variant were administered with a single IP dose of either saline (n=16) or Ronapreve, 400 μg/mouse, in saline (n=16). Data for individual animals are shown with the mean value represented by a black line. NS, not significant; *, P ≤0.05 (unpaired, two-tailed t-test).

In contrast, in the Omicron infected mice the amount of sgE RNA detected in the nasal turbinates was only marginally reduced at both 4 dpi (log10 fold decrease: −0.243, P=0.267) and 6 dpi (log10 fold decrease: −0.065, P=0.973) in the Ronapreve treated mice compared to the saline controls. The same effect was observed in the lung at 4 dpi (log10 fold reduction: −0.312, P=0.149) whereas it was similar at 6 dpi (log10 fold increase: 0.130, P=0.390). The results highlight the diminished *in vivo* antiviral potency of Ronapreve against the Omicron variant.

### Differences in viral replication between Delta and Omicron variants

Mice were challenged with a comparable amount of virus (10^3^ PFU) of both SARS-CoV-2 variants. However, comparison of the sgE RNA levels in the tissues, of the saline treated animals at both time points, showed that infection with the Omicron variant generally yielded lower viral loads. In the nasal turbinate samples, a log10 fold lower viral RNA level of −0.243, P=0.267 (4 dpi) and −2.043, P=0.099 (6 dpi) was observed in the Omicron group. In the lung, a log10 fold lower viral RNA level of −0.353, P=0.137 (4 dpi) and −0.561, P=0.085 (6 dpi) was observed. Detailed information on viral loads in individual animals is provided in Supplemental Table S1. Similar trends have been reported in Omicron infected mice displaying a lower viral load in both upper and lower respiratory tracts.^33^

### The effect of Ronapreve on pulmonary changes and viral spread to the brain after infection with the Delta and Omicron variants

At 4 dpi, productive virus infection was also confirmed by immunohistochemistry in all groups of infected mice. In saline treated animals, virus antigen was detected in epithelial cells in the nasal mucosa in all Delta infected mice, but not in Omicron infected mice. This trend was consistent with nasal turbinate PCR data with lower sgE RNA levels in the saline treated Omicron infected mice compared to the Delta infected mice. Viral antigen was detected in the lung of 5 of the 6 Delta infected mice. Infection was generally widespread and represented by numerous large, partly coalescing patches of alveoli with positive type I and type II pneumocytes (Supplementary Figure S1a). Histologically, this was accompanied by the presence of activated type II pneumocytes, occasional syncytial cells and degenerate alveolar epithelial cells as well as scattered desquamed cells within alveolar lumina and increased interstitial cellularity; mild vasculitis was also seen. Also in the Omicron infected mice, viral antigen was detected in the lung (7 of 8 animals). Expression was seen in disseminated small patches of alveoli with positive pneumocytes (Supplementary Figure S1b) and was overall less abundant than in the Delta infected mice, confirming the virological results. Infection accompanied by focal areas with increased interstitial cellularity and desquamation of a few alveolar cells, some infiltrating lymphocytes and macrophages as well as occasional mild vasculitis. The lung PCR data revealed no significant difference between the saline treated Omicron infected mice and the Delta infected mice (P=0.137). Interestingly, animal C2.5 (Table S1) was negative for both viral antigen and sgE RNA, demonstrating consistency between the immunohistochemistry and PCR data.

After Ronapreve treatment, at 4 dpi, virus antigen was detected in epithelial cells in the nasal mucosa in 4 of the 8 Delta infected mice, and in 1 of the 8 Omicron infected mice. Viral antigen was detected in the lung of 7 of the 8 Delta infected mice. It was generally less extensively expressed than in the saline treated group and seen in the pneumocytes of small disseminated patches of alveoli (Supplementary Figure S1c). Infection was accompanied by similar histological changes as in the untreated mice, but these were less extensive. In Omicron infected mice, viral antigen was detected in 5 of the 8 animals, with a similar extent and distribution as the Delta infected mice and the untreated Omicron infected group alveoli (Supplementary Figure S1d), and with histological changes similar to those seen in the untreated Omicron infected mice in nature and extent.

At 6 dpi, productive virus infection was still observed in all groups of infected mice. In saline treated animals, virus antigen was detected in epithelial cells in the nasal mucosa in all Delta infected mice but in none of the Omicron infected mice. Lower sgE RNA levels in the saline treated Omicron infected mice compared to the Delta infected mice was observed, albeit not statistically significant (P=0.099). Examination of the lungs revealed viral antigen in the lungs of 4 of the 6 Delta infected animals, mainly in numerous, often large disseminated patches of alveoli (Supplementary Figure S1e), and most intense in association with large focal areas of increased interstitial cellularity that contained activated type II pneumocytes, occasional syncytial cells and degenerate and/or desquamed alveolar epithelial cells. In Omicron infected mice, viral antigen expression was detected in all 8 animals, generally in numerous disseminated, mainly small patches of alveoli (Supplementary Figure S1f). It was overall less extensive than in the Delta infected animals at this time point, further supporting the virology results, with lower sgE RNA levels in the saline treated Omicron infected mice compared to the Delta infected mice, albeit not statistically significant (P=0.085). Infection was accompanied by mild histological changes, represented by small focal areas with desquamed alveolar epithelial cells and mild mononuclear infiltration.

After Ronapreve treatment, at 6 dpi, a different histopathological picture was observed in the Delta infected mice. When viral antigen was detected (7/8 mice), its expression was very limited (Supplementary Figure S1g). Histologically, multifocal small, delineated, dense parenchymal mononuclear infiltrates were found (Fig. 3). These were comprised of macrophages (Iba1+), with lesser T cells (CD3+) and B cells (CD45R+) and also seen to involve vessels, where a patchy vasculitis, with focal infiltration of the vascular wall, stretching into a focal perivascular infiltrate, was observed (Fig. 3). These lesions often contained a few infected alveolar epithelial cells and some free viral antigen, consistent with debris of infected cells (Fig. 3f). Otherwise, viral antigen expression was limited to epithelial cells in a few small patches of alveoli (Supplementary Figure S1g). All 7 Omicron infected animals were found to harbor viral antigen in the lung (Supplementary Figure S1h), the extent and distribution of viral antigen was similar to that at 4 dpi, seen as disseminated small patches of alveoli with positive pneumocytes. The accompanying histological changes were generally mild and as described for the control mice, though focal infiltrates similar to those seen in the Delta infected treated mice were also seen, albeit overall less pronounced and less delineated. Detailed information on histological findings, viral antigen expression and viral loads in individual animals is provided in Supplemental Table S1.

**Figure 3.**
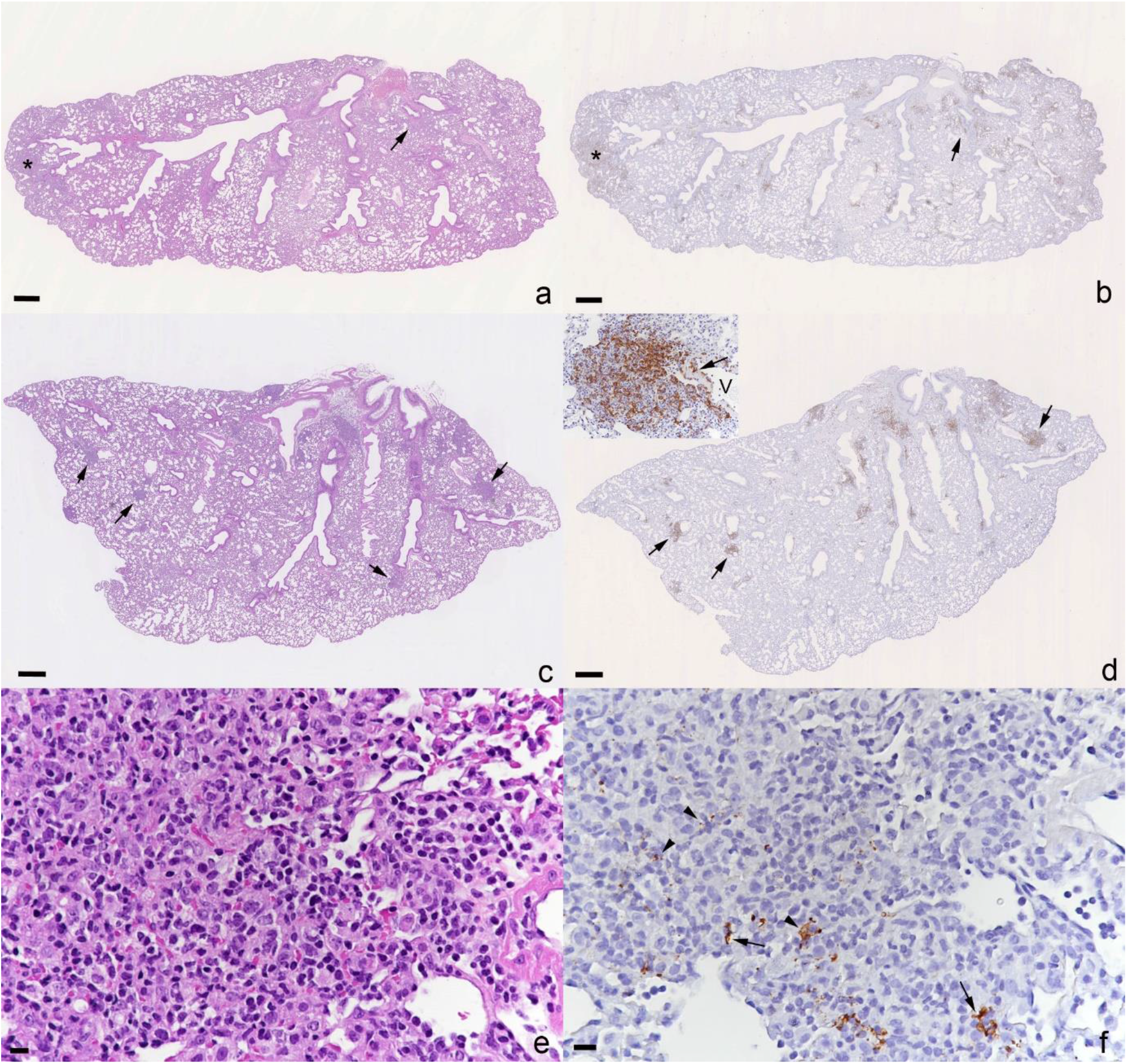
Lungs, K18-hACE2 mice at day 6 post infection with 10^3^ PFU of SARS-CoV-2 Delta variant (B.1.617.2), followed after 24 hours by an intraperitoneal injection of 100 μL saline control or 400 μg Ronapreve, diluted in saline. **a, b)** Saline treated animal (C3.1). The parenchyma shows a focal consolidated area (asterisk) and several areas with increased cellularity (arrow) in the parenchyma in which macrophages (Iba1+) are the dominant inflammatory cells (a: HE stain; b: Iba1 immunohistology; bars = 500 μm). **c-f)** Ronapreve treated animal (R3.2). **c, d)** The parenchyma exhibits several well delineated dense inflammatory infiltrates (arrows) that are dominated by macrophages (Iba1+). Inset (d): Closer view of a focal inflammatory infiltrate. Macrophages are the dominant cells and are also seen to emigrate from a vessel (V; arrow). (c: HE stain; d: Iba1 immunohistology; bars = 500 μm). **e, f)** Closer view of a focal inflammatory infiltrate. Viral antigen is found within a few pneumocytes (arrows) and cell free or phagocytosed within macrophages (arrowheads). Bars = 25 μm.

**Figure 4.**
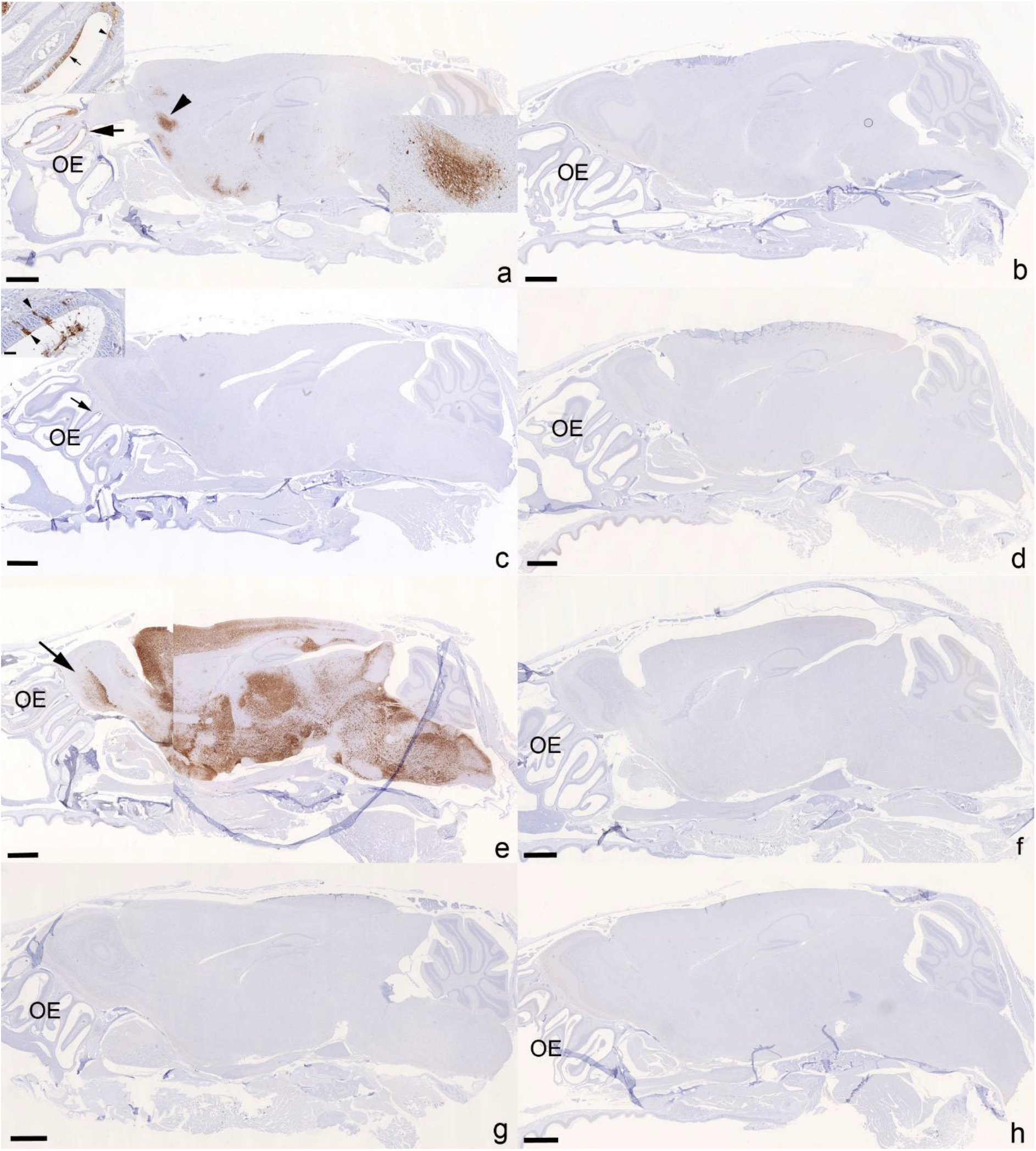
Heads with olfactory epithelium (OE) and brain, K18-hACE2 mice. SARS-CoV-2 N expression at day 4 (a-d) and day 6 (e-h) post infection with 10^3^ PFU of SARS-CoV-2 Delta variant (B.1.617.2; a, c, e, g) or Omicron variant (b, d, f, h), followed after 24 hours by an intraperitoneal injection of 100 μL saline control (a, b, e, f) or 400 μg Ronapreve (c, d, g, h), diluted in saline. **a)** Delta variant infected mouse (C1.1) treated with saline control, 4 dpi. The virus is widespread in the OE (arrow and inset showing a large patch of positive epithelial cells (arrow) and a few individual positive epithelial cells (arrowhead)) and has spread to the brain; there are patches of neurons positive for viral antigen, in frontal cortex, cerebral nucei (caudoputamen), hypothalamus/thalamus, midbrain and pons. The arrowhead depicts a large patch of positive neurons in the frontal cortex of which a closer view is provided in the inset. **b)** Omicron variant infected mouse (C2.3) treated with saline control, 4 dpi. There is no evidence of viral antigen expression in the OE and the brain. **c)** Delta variant infected mouse (R1.1) treated with Ronapreve, 4 dpi. There is no evidence of viral antigen expression in the brain. The OE exhibits a small patch with positive epithelial cells. Inset: OE with viral antigen expression in intact individual olfactory epithelial cells (arrowheads) and in degenerate cells in the lumen of the nasal cavity. **d)** Omicron variant infected mouse (R2.5) treated with Ronapreve, 4 dpi. There is no evidence of viral antigen expression in the OE and the brain. **e)** Delta variant infected mouse (C3.3) treated with saline control, 6 dpi. There is widespread viral antigen expression in abundant neurons throughout the brain including the olfactory bulb (left, arrow), with the exception of the cerebellum. **f)** Omicron variant infected mouse (C4.1) treated with saline control, 6 dpi. There is no evidence of viral antigen expression in the OE and the brain. **g)** Delta variant infected mouse (R3.3) treated with Ronapreve, 6 dpi. There is no evidence of viral antigen expression in the OE and the brain. **h)** Omicron variant infected mouse (C4.7) treated with Ronapreve, 6 dpi. here is no evidence of viral antigen expression in the OE and the brain. Immunohistology, hematoxylin counterstain, and HE stain (insets). Bars = 1 mm.

We and others have previously shown that wildtype and VOC SARS-CoV-2s readily spread to the brain in K18-hACEs mice; Omicron variants appear not to have the same effect, remaining unaltered and without viral antigen expression^32^. The current study confirmed these findings. Brain infection was a rather consistent finding in the untreated Delta infected animals. At 4 dpi, viral antigen was detected multifocally in neurons in the brain in 4 of the 6 mice (Supplementary Fig S2a), all exhibiting viral antigen also in the nasal mucosa, and in particular in the olfactory epithelium. Viral antigen was also detected in nerve fibres or a variable amount of neurons in the olfactory bulb, consistent with viral spread from the nasal mucosa, via the olfactory plate.^34^ There was no evidence of an inflammatory response. In contrast, none of the Omicron infected animals were found to harbor viral antigen in the nasal mucosa and the brain (Supplemental Fig 2b). At 6 dpi, brain infection was confirmed by immunohistology in 5 of the 6 Delta infected mice; viral antigen expression was still restricted to neurons but was generally very widespread (Supplementary Fig S2e); in two mice it was accompanied by mild perivascular mononuclear infiltrates consistent with a non-suppurative encephalitis. The nasal mucosa, and in particular the olfactory epithelium, often also underlying nerve fibres were found to harbor viral antigen also at this stage. Again, the Omicron infected mice did not show any viral antigen expression in nasal mucosa and brain (Supplementary Fig S2f).

After Ronapreve treatment, there was no evidence of viral antigen expression in the brain of any Delta infected animal at 4 and 6 dpi (each n=8; Supplementary Fig 2c and g); however, the nasal mucosa harbored infected epithelial cells at both time points, in 4 of the 8 animals at 4 dpi, and in 6 of the 8 animals at 6 dpi. Omicron infected, Ronapreve treated animals were equally negative for viral antigen in the brain and the nasal mucosa at both time points, with the exception of one mouse at 4 dpi. Detailed information on histological findings and viral antigen expression in the brains of individual animals is provided in Supplemental Table S1.

## Figure legends

**Supplemental Figure S1.**
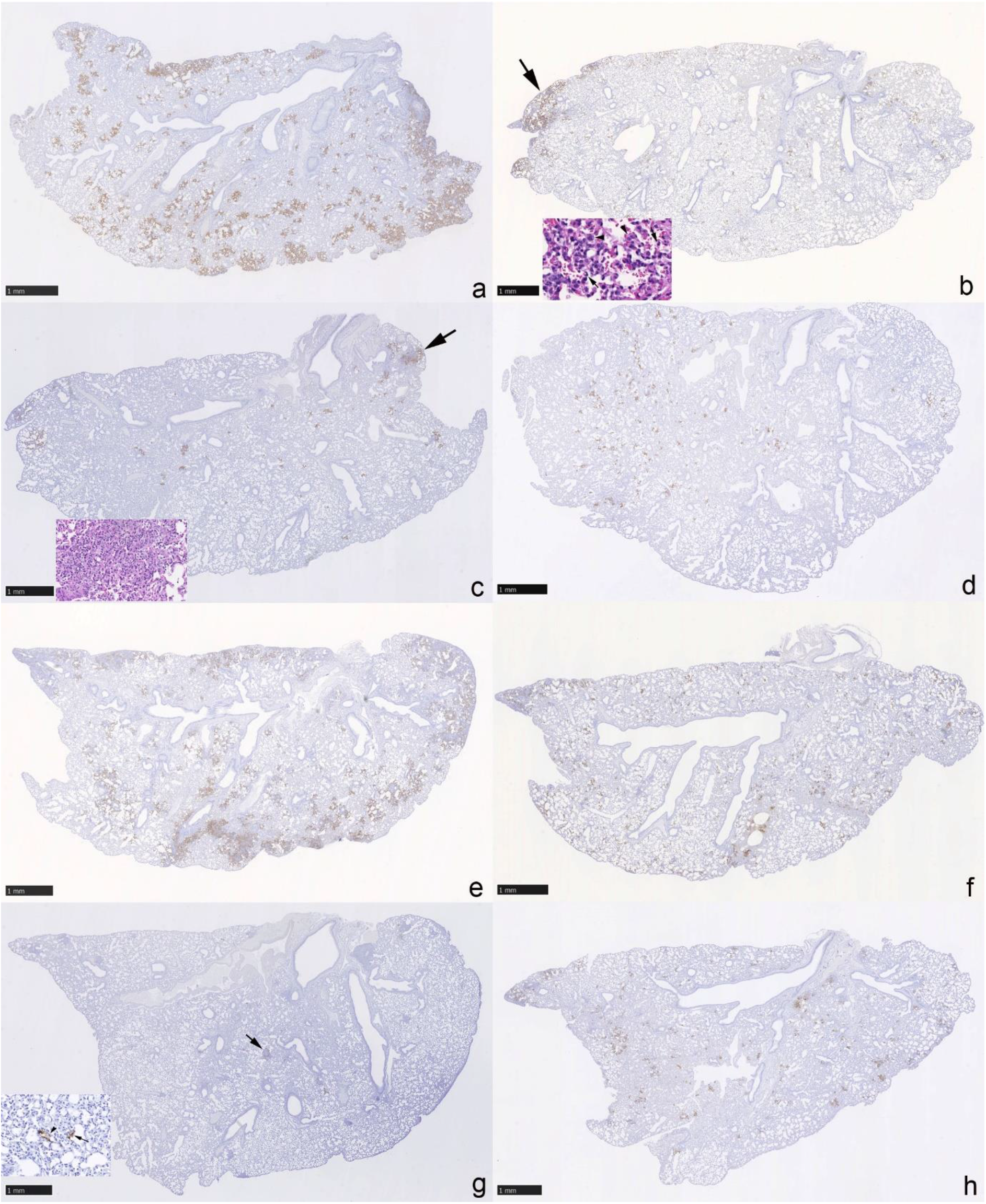
Left lung, longitudinal sections, K18-hACE2 mice. SARS-CoV-2 N expression at day 4 (a-d) and day 6 (e-h) post infection with 10^3^ PFU of SARS-CoV-2 Delta variant (B.1.617.2; a, c, e, g) or Omicron variant (b, d, f, h), followed after 24 hours by an intraperitoneal injection of 100 μL saline control (a, b, e, f) or 400 μg Ronapreve (c, d, g, h), diluted in saline. **a)** Delta variant infected mouse (C1.2) treated with saline control, 4 dpi. Abundant large, partly coalescing patches of alveoli with positive epithelial cells are found disseminated throughout the parenchyma. **b)** Omicron variant infected mouse (C2.1) treated with saline control, 4 dpi. There are multiple disseminated small patches of alveoli with positive epithelial cells. A large patch (arrow) of positive alveoli is seen in association with focal desquamation of alveolar epithelial cells (inset: arrows) and the presence of activated and syncytial type II pneumocytes (inset: arrowheads). **c)** Delta variant infected mouse (R1.5) treated with Ronapreve, 4 dpi. There are numerous small disseminated patches of alveoli with positive epithelial cells, and larger patches (arrow) in association with focal activation and syncytia formation in type II pneumocytes, desquamation of alveolar epithelial cells, occasional degenerate cells and a few infiltrating lymphocytes and neutrophils (inset). **d)** Omicron variant infected mouse (R2.6) treated with Ronapreve, 4 dpi. Viral antigen expression is seen in epithelial cells of random small patches of alveoli. **e)** Delta variant infected mouse (C3.5) treated with saline control, 6 dpi. Multifocal extensive, partly coalescing large patches of alveoli with positive epithelial cells are found disseminated throughout the parenchyma. **f)** Omicron variant infected mouse (C4.8) treated with saline control, 6 dpi. There are multiple disseminated, mainly small patches of alveoli with pos epithelial cells. **g)** Delta variant infected mouse (R3.4) treated with Ronapreve, 6 dpi. There are disseminated very small patches of alveoli with positive epithelial cells (inset). Positive cells are also observed in focal infiltrates (arrow; see Fig. 3). **h)** Omicron variant infected mouse (C4.8) treated with Ronapreve, 6 dpi. There are numerous disseminated, mainly small patches of alveoli with pos epithelial cells. Immunohistology, hematoxylin counterstain, and HE stain (insets). Bars = 1 mm.

## Discussion

The current study made use of a reliable animal model of SARS-CoV-2 infection to confirm and characterize the effect of Ronapreve on established infections with the Delta variant and confirm its ineffectiveness for BA.1 Omicron infections. Indeed, it provides increased certainty in the absence of effect for Ronapreve which complements *in vitro* neutralisation data for this variant.^23, 35^ However, it confirms efficacy for the Delta variant and provides evidence that monoclonal antibodies might limit the virus spread into the brain when deployed against susceptible variants. In addition, it reveals pathological processes that can develop in the lungs when Ronapreve is applied after the Delta variant has reached the lungs.

A very rapid comparable decline in body weight was observed in all groups of mice, at 2 dpi, different from previous studies that showed consistent weight drop only at 3 dpi. This early drop is likely a response to the invasiveness and additional handling associated with the intraperitoneal dosing. The weight gain towards day 3 moved body weights to levels observed in a previous study in which K18-hACE2 transgenic mice were infected with the same virus variants at the same dose, but were not treated any further.^33^ By the end of the study, all but the saline-control animals infected with the Delta variant had regained more weight. This observation is in agreement with the authors’ previous evaluation of the pathogenicity of these variants in K18 hACE2 transgenic mice.^33^ Mice infected with the Omicron variant generally carried less sub-genomic viral RNA than Delta variant-infected mice in both nasal turbinates and lungs at 4 and 6 dpi, which is consistent with previous reports and lower viral replication of Omicron in the respiratory tract and lungs at these time points.^33^ The histological and immunohistological results support this finding. At both time points, viral antigen expression was less widespread after Omicron infection, and the histological changes, representing focal areas of alveolar damage with associated inflammatory response, were less severe.

Consistent with the clinical evidence through body weight measurements, levels of sub-genomic RNA were reduced in both nasal turbinates and lung of mice infected with the Delta variant after Ronapreve treatment compared to controls, at both time points. This finding was complemented by the results of the histological and immunohistological examinations. While the control group at 4 dpi exhibited, in the majority, multifocal areas with activation and syncytia formation of type II pneumocytes, some desquamation of alveolar epithelial cells and occasional vasculitis, with extensive multifocal viral antigen expression, the lungs of the Ronapreve treated mice were either found unaltered and free of viral antigen, or exhibited a few small focal areas with alveolar damage and small patches of infected alveoli, but no evidence of vasculitis. This suggests that post-exposure Ronapreve treatment reduces pulmonary damage. Two days later, the difference between saline control and Ronapreve treated mice was even greater. In the former, the lesions observed at 4 dpi were found to persist and the accompanying inflammatory response had intensified, resulting in larger consolidated areas and perivascular leukocyte infiltrates, with extensive multifocal viral antigen expression in large patches of alveoli. After Ronapreve treatment, a different type and extent of inflammation and viral antigen expression was observed. Viral antigen was only seen in a few very small patches of alveolar epithelial cells and within small, delineated focal macrophage dominated, i.e. granulomatous parenchymal infiltrates that also harbored viral antigen. Their proximity to and frequent continuity with identical focal infiltrates of vascular walls and the presence of viral antigen not only within pneumocytes but also within macrophages in these lesions indicate that they result from focal recruitment of macrophages into the parenchyma in response to virus. Ronapreve represents human antibodies that target the spike protein on the surface of SARS-CoV-2. After a single application, the antibodies will not have induced an immune response in the mice, instead they will likely have bound to the Fc receptors of the murine macrophages.^36^ Considering that the granulomatous reaction was not observed in the saline controls, it is likely that it represents the local response to antibody-opsonised virus that is phagocytosed by macrophages. A previous study that histologically examined the lungs of mice treated with a neutralizing antibody at 2 dpi as late as 21 days post infection found unaltered lungs with only scarce lymphoid aggregates,^27^ indicating that these local processes can dissolve with time. Murine models are generally robust for identifying potential pathological effects of therapeutic interventions. The implications of these findings to clinical deployment of monoclonal antibodies is currently uncertain, but further robust assessment incorporating morphological measures in parallel to virological measures is warranted.

The immunohistological examination also revealed a further positive effect of the Ronapreve treatment. As expected, the Delta variant had already spread to the brain in some animals by day 4 and was found widespread in the brain at 6 dpi in mice that had received the saline control.^32^ At the later time point it had induced a mild inflammatory response in some animals. After Ronapreve treatment, there was no evidence of viral antigen expression in the brain, and no inflammatory change. These findings suggest that Ronapreve treatment post exposure might inhibit viral spread into the brain. Whether infection of the brain is completely blocked or only substantially reduced, requires further investigations, particularly at the molecular level. In light of previous studies which showed that, in K18-hACE2 mice, the virus reaches the brain predominantly via the olfactory route^32, 34^ and considering Ronapreve treatment reduces viral loads in the nasal turbinates it is probable that Ronapreve inhibits brain infection by reducing the risk of virus spread from the olfactory epithelium to the underlying nerves, then the olfactory bulb and into the brain.

Conversely, no significant impact of Ronapreve on sub-genomic RNA over 6 days was observed in mice infected with BA.1 Omicron, which is consistent with a loss of neutralisation of this variant. The doses used in the current study were 2-fold higher than those for which virological efficacy was demonstrated in K18-hACE2 transgenic mice previously for other variants^22^ which reinforces the conclusion that activity against Omicron is ablated for Ronapreve. The immunohistological results further support the virological findings, as they indicate no or only mild reduction of viral antigen expression after Ronapreve treatment. At 6 dpi, a limited granulomatous response was observed in some lungs, suggesting some virus opsonization, though only to a very low extent.

Curiously, the magnitude of the reduction in Delta sub-genomic RNA was lower in the present study than that reported for total RNA in the previous study despite the higher dose.^22^ At the time of writing, the authors are unaware of other studies that have investigated the efficacy of Ronapreve for Delta in this model, but neutralisation of Delta was not meaningfully compromised *in vitro.^13^* Differences in the endpoint (sub-genomic versus total RNA measurements) make it difficult to draw firm conclusions from these observations but underscore the importance of *in vivo* evaluation of the efficacy of interventions against new and future variants.

The experimental design employed here reflects treatment whereby the intervention was applied subsequent to the inoculation of the animals with virus. Several other studies that have sought to assess continued efficacy of monoclonal antibodies against later Omicron sub-lineages have utilized prophylactic designs where the antibody is administered prior to inoculation of the animals with virus ^37–39^. Because of the differences in viral load when the intervention is introduced, it is well established for antiviral interventions that the bar is much higher to achieve efficacy in treatment than it is for prophylaxis. The data presented here clearly demonstrate that *in vivo* designs reflecting the intended treatment use case are achievable and demonstrate efficacy for monoclonal antibodies against susceptible variants. Extreme caution should be taken when interpreting *in vivo* data from prophylactic designs when making an assessment of the likely continued efficacy in treatment, and where animal data are used to support candidacy of interventions, *in vivo* studies should be designed to be reflective of the intended use case in humans.

A limitation of the current study is that serum concentrations of the Ronapreve antibodies were not measured in order to facilitate a comparison with exposures observed in humans. However, the lack of virological efficacy for BA.1 Omicron despite a demonstrable impact upon Delta, coupled with the higher doses used here compared with a previous study with earlier variants^22^ allows for confidence in the outcome despite this deficit.

## Acknowledgements

The authors are grateful to the technical staff at the Histology Laboratory, Institute of Veterinary Pathology, Vetsuisse Faculty, University of Zurich, for excellent technical support.

## Transparency declarations

AO and SR are Directors of Tandem Nano Ltd and co-inventors of patents relating to drug delivery. AO has been co-investigator on funding received by the University of Liverpool from ViiV Healthcare and Gilead Sciences unrelated to COVID-19 in the past 3 years. AO has received personal fees from Gilead and Assembly Biosciences in the past 3 years unrelated to COVID-19. AO was a member of the Trial Management Group for the AGILE phase I/II platform trial until January 2023 and AGILE received funding from Ridgeback and GSK in the past 3 years for which AO was not a co-investigator. SR has received research funding from ViiV and AstraZeneca and consultancy from Gilead not related to the current paper. No other conflicts are declared by the authors.

