## Supplementary Table S1. for "Ronapreve (REGN-CoV; casirivimab and imdevimab) reduces the viral burden and alters the pulmonary response to the SARS-CoV-2 Delta variant (B.1.617.2) in K18-hACE2 mice using an experimental design reflective of a treatment use case"

**Supplementary Table S1.** Relevant histological changes and SARS-CoV-2 nucleoprotein expression in K18-hACE2 mice infected with SARS-CoV-2 Delta or Omicron BA.1 variant at an infectious dose of $10^3$ PFU intranasally, treated 24 hours post infection with saline (controls) or Ronapreve, and euthanised at 4 or 6 days post infection.

| Animal No | Infection, dpi[1] | Histological changes and viral antigen expression | Virology (PCR)[2] |
|---|---|---|---|
| **Saline treated (control) animals** | | | |
| C1.1 | Delta, 4 dpi | **Nasal mucosa[3] (HE)**: abundant deg EC, abundant leukocytes and deg cells in lumen<br>**vAg**: individual and patches of intact and deg pos REC and OEC, pos deg cells in lumen | 54585 |
| | | **Lung (HE)**: multifocal to disseminated activated type II pc, occ syncytial cells and deg AEC, scattered desquamed AEC; focal IIC; occ vasculitis<br>**vAg**: abundant large, partly coalescing disseminated patches of alveoli with pos AEC | 2981935 |
| | | **Brain (HE)**: NHA<br>**vAg**: patches of several pos neurons (frontal cortex, brainstem, hippocampus) | N/A |
| C1.2 | Delta, 4 dpi | **Nasal mucosa (HE)**: NHA<br>**vAg**: individual and patches of intact pos OEC | 486818 |
| | | **Lung (HE)**: multifocal to disseminated activated type II pc, occ syncytial cells and deg AEC, scattered desquamed AEC; focal IIC; occ vasculitis<br>**vAg**: abundant large, partly coalescing disseminated patches of alveoli with pos AEC | 4023006 |
| | | **Brain (HE)**: NHA<br>**vAg**: patches with variable number of pos neurons (olfactory bulb, frontal cortex, brainstem, hippocampus, medulla oblongata) | N/A |
| C1.3 | Delta, 4 dpi | **Nasal mucosa (HE)**: several deg OEC<br>**vAg**: individual and patches of intact pos OEC | 395496 |
| | | **Lung (HE)**: NHA<br>**vAg**: neg | 4156 |
| | | **Brain (HE)**: NHA<br>**vAg**: patches with variable number of pos neurons (olfactory bulb, frontal cortex, brainstem, hippocampus, medulla oblongata) | N/A |
| C1.4 | Delta, 4 dpi | **Nasal mucosa (HE)**: a few deg EC<br>**vAg**: a few individual pos intact and deg REC and OEC | 886299 |
| | | **Lung (HE)**: multifocal to disseminated activated type II pc, occ syncytial cells and deg AEC, a few desquamed AEC, focal IIC; occ vasculitis<br>**vAg**: abundant large, partly coalescing disseminated patches of alveoli with pos AEC (particularly intense in affected areas) | 9142064 |
| | | **Brain (HE)**: NHA<br>**vAg**: neg | N/A |
| C1.5 | Delta, 4 dpi | **Nasal mucosa (HE)**: some deg OEC<br>**vAg**: several and large patches of pos intact and deg OEC | 1111073 |
| | | **Lung (HE)**: multifocal areas with activated type II pc, occ syncytial cells, focal IIC, minimal AEC desquamation<br>**vAg**: several disseminated, variably sized patches of alveoli with pos AEC | 2564 |
| | | **Brain (HE)**: NHA<br>**vAg**: a few individual pos neurons in olfactory bulb and frontal cortex | N/A |
| C1.6 | Delta, 4 dpi | **Nasal mucosa (HE)**: some deg EC<br>**vAg**: patches of pos intact and deg REC and OEC | 500837 |
| | | **Lung (HE)**: multifocal areas with activated type II pc, occ syncytial cells and deg AEC, a few desquamed AEC within alveolar lumina and IIC; occ vasculitis<br>**vAg**: abundant large, partly coalescing disseminated patches of alveoli with pos AEC | 5950179 |
| | | **Brain (HE)**: NHA<br>**vAg**: neg | N/A |
| C2.1 | Omicron, 4 dpi | **Nasal mucosa (HE)**: NHA<br>**vAg**: neg | 1570 |
| | | **Lung (HE)**: a few focal subpleural areas of IIC and AEC desquamation, with type II pc activation and some LC, NL and macrophages; mild pv and pb LC infiltration<br>**vAg**: multiple disseminated small patches of alveoli with pos AEC, large patch in association with area of AEC desquamation | 2715791 |
| | | **Brain (HE)**: NHA<br>**vAg**: neg | N/A |
| C2.2 | | **Nasal mucosa (HE)**: NHA<br>**vAg**: neg | 10117 |

| | | | |
|---|---|---|---|
| | Omicron, 4 dpi | **Lung (HE)**: small focal areas of IIC and AEC desquamation, with some LC and macrophages<br>**vAg**: several disseminated small patches of alveoli with pos AEC | 990308 |
| | | **Brain (HE)**: NHA<br>**vAg**: neg | N/A |
| C2.3 | Omicron, 4 dpi | **Nasal mucosa (HE)**: NHA<br>**vAg**: neg | 9174 |
| | | **Lung (HE)**: a few focal areas of IIC, activated type II pc, some LC and macrophages; a few vessels with mild vasculitis<br>**vAg**: several disseminated small patches of alveoli with pos AEC | 1630021 |
| | | **Brain (HE)**: NHA<br>**vAg**: neg | N/A |
| C2.4 | Omicron, 4 dpi | **Nasal mucosa (HE)**: NHA<br>**vAg**: neg | 6483 |
| | | **Lung (HE)**: a few focal areas of IIC; a few vessels with mild pv mononuclear infiltration; one large bronchiole with a few deg BEC<br>**vAg**: several disseminated small patches of alveoli with pos AEC; one large bronchiole with several pos, partly deg BEC | 2238181 |
| | | **Brain (HE)**: NHA<br>**vAg**: neg | N/A |
| C2.5 | Omicron, 4 dpi | **Nasal mucosa (HE)**: NHA<br>**vAg**: neg | 2129 |
| | | **Lung (HE)**: NHA<br>**vAg**: neg | <LOD |
| | | **Brain (HE)**: NHA<br>**vAg**: neg | N/A |
| C2.6 | Omicron, 4 dpi | **Nasal mucosa (HE)**: NHA<br>**vAg**: neg | 3633 |
| | | **Lung (HE)**: several small focal areas of pv mononuclear (macrophages, LC, a few deg cells) infiltration with vasculitis; focal subpleural area with IIC and mild AEC desquamation<br>**vAg**: several disseminated small patches of alveoli with pos AEC; larger area in association with AEC desquamation | 2463208 |
| | | **Brain (HE)**: NHA<br>**vAg**: neg | N/A |
| C2.7 | Omicron, 4 dpi | **Nasal mucosa (HE)**: NHA<br>**vAg**: neg | 5157 |
| | | **Lung (HE)**: a few small focal areas of (pv) mononuclear (macrophages, LC, a few deg cells) infiltrates with vasculitis<br>**vAg**: numerous disseminated, mainly small patches of alveoli with pos AEC | 1175908 |
| | | **Brain (HE)**: NHA<br>**vAg**: neg | N/A |
| C2.8 | Omicron, 4 dpi | **Nasal mucosa (HE)**: NHA<br>**vAg**: neg | 12632 |
| | | **Lung (HE)**: several small focal areas of pv mononuclear (macrophages, LC, a few deg cells) infiltration with vasculitis; focal subpleural area with IIC and some AEC desquamation<br>**vAg**: numerous disseminated, mainly small patches of alveoli with pos AEC; larger area in association with AEC desquamation | 1849172 |
| | | **Brain (HE)**: NHA<br>**vAg**: neg | N/A |
| C3.1 | Delta, 6 dpi | **Nasal mucosa (HE)**: NHA<br>**vAg**: individual and patches of intact pos OEC | 64788 |
| | | **Lung (HE)**: mild to moderate, almost diffuse IIC; multifocal areas with activated type II pc, occ syncytial cells and deg AEC, occ AEC desquamation; occ vasculitis and pv leukocyte infiltrates<br>**vAg**: numerous variably sized disseminated patches of alveoli with pos AEC; most intense in areas with AEC desquamation | 6146147 |
| | | **Brain (HE)**: very mild pv leukocyte infiltration affecting a few vessels in brain stem<br>**vAg**: very numerous pos neurons throughout entire brain (except for cerebellar cortex) | N/A |
| | | **Nasal mucosa (HE)**: NHA | 1313784 |

| | | | |
|---|---|---|---|
| C3.2 | Delta, 6 dpi | **vAg**: individual and patches of intact pos OEC | |
| | | **Lung (HE)**: multifocal areas with activated type II pc, occ syncytial cells and deg AEC, macrophages and LC, occ AEC desquamation<br>**vAg**: numerous disseminated large patches of alveoli with pos AEC; most intense in areas with AEC desquamation | 7906870 |
| | | **Brain (HE)**: NHA<br>**vAg**: patches of and individual pos neurons throughout entire brain (except for cerebellar cortex) | N/A |
| C3.3 | Delta, 6 dpi | **Nasal mucosa (HE)**: a few deg OEC<br>**vAg**: individual and patches of intact and deg pos OEC | 389193 |
| | | **Lung (HE)**: focal area with activated type II pc, occ deg AEC, infiltrating interstitial NL, macrophages and LC, occ desquamed AEC<br>**vAg**: one large patch of alveoli with pos AEC close to focal lesion | 464823 |
| | | **Brain (HE)**: NHA<br>**vAg**: numerous pos neurons throughout entire brain (except for cerebellar cortex) | N/A |
| C3.4 | Delta, 6 dpi | **Nasal mucosa (HE)**: a few deg OEC<br>**vAg**: individual and patches of intact pos OEC | 104123 |
| | | **Lung (HE)**: NHA<br>**vAg**: neg | 19343 |
| | | **Brain (HE)**: very mild pv leukocyte infiltration of a few vessels in brain stem and adjacent leptomeninx<br>**vAg**: numerous pos neurons throughout entire brain (except for cerebellar cortex) | N/A |
| C3.5 | Delta, 6 dpi | **Nasal mucosa (HE)**: occ deg OEC<br>**vAg**: with several pos intact and deg OEC | 41230 |
| | | **Lung (HE)**: large focal consolidated areas with activated type II pc, deg cells, macrophages and LC; other large areas with a few desquamed AEC, activated type II pc, a few macrophages, LC and NL; mild pv mononuclear infiltrates<br>**vAg**: multifocal extensive, partly coalescing large patches of alveoli with pos AEC, also adjacent to consolidated areas | 5691395 |
| | | **Brain (HE)**: NHA<br>**vAg**: neg | N/A |
| C3.6 | Delta, 6 dpi | **Nasal mucosa (HE)**: NHA<br>**vAg**: rare pos intact OEC | 57656 |
| | | **Lung (HE)**: NHA<br>**vAg**: neg | 2013 |
| | | **Brain (HE)**: NHA<br>**vAg**: patches of pos neurons in frontal cortex, brainstem and medulla oblongata | N/A |
| C4.1 | Omicron, 6 dpi | **Nasal mucosa (HE)**: NHA<br>**vAg**: neg | 3619 |
| | | **Lung (HE)**: two small focal pv mononuclear (macrophages, LC, a few deg cells) infiltrates with vasculitis<br>**vAg**: a few small patches of alveoli with pos AEC, a few pos cells in focal infiltrate | <LOD |
| | | **Brain (HE)**: NHA<br>**vAg**: neg | N/A |
| C4.2 | Omicron, 6 dpi | **Nasal mucosa (HE)**: NHA<br>**vAg**: neg | 2187 |
| | | **Lung (HE)**: several small focal areas of pv mononuclear (macrophages, LC, a few deg cells) infiltrates with vasculitis; small focal areas with AEC desquamation<br>**vAg**: multiple disseminated, mainly small patches of alveoli with pos AEC, larger area in association with AEC desquamation | 1563149 |
| | | **Brain (HE)**: NHA<br>**vAg**: neg | N/A |
| C4.3 | Omicron, 6 dpi | **Nasal mucosa (HE)**: NHA<br>**vAg**: neg | 6795 |
| | | **Lung (HE)**: several small focal areas of pv mononuclear (macrophages, LC, a few deg cells) infiltrates with vasculitis; small focal areas with AEC desquamation<br>**vAg**: multiple disseminated, mainly small patches of alveoli with pos AEC, larger area in association with AEC desquamation | 65806 |
| | | **Brain (HE)**: NHA<br>**vAg**: neg | N/A |
| C4.4 | | **Nasal mucosa (HE)**: NHA<br>**vAg**: neg | 6057 |

| | | | |
|---|---|---|---|
| | Omicron, 6 dpi | **Lung (HE)**: several small focal areas of pv mononuclear (macrophages, LC, a few deg cells) infiltrates with vasculitis; small focal areas with AEC desquamation<br>**vAg**: multiple disseminated, mainly small patches of alveoli with pos AEC, slightly larger area in association with AEC desquamation | 1071043 |
| | | **Brain (HE)**: NHA<br>**vAg**: neg | N/A |
| C4.5 | Omicron, 6 dpi | **Nasal mucosa (HE)**: NHA<br>**vAg**: neg | <LOD |
| | | **Lung (HE)**: several small focal areas of (pv) mononuclear (macrophages, LC, a few deg cells) infiltrates with vasculitis; small focal areas with AEC desquamation<br>**vAg**: numerous random small patches of alveoli with pos AEC | 2066925 |
| | | **Brain (HE)**: NHA<br>**vAg**: neg | N/A |
| C4.6 | Omicron, 6 dpi | **Nasal mucosa (HE)**: NHA<br>**vAg**: neg | 1352 |
| | | **Lung (HE)**: several small focal areas of (pv) mononuclear (macrophages, LC, a few deg cells) infiltrates with vasculitis<br>**vAg**: many random small patches of alveoli with pos AEC | 1865896 |
| | | **Brain (HE)**: NHA<br>**vAg**: neg | N/A |
| C4.7 | Omicron, 6 dpi | **Nasal mucosa (HE)**: NHA<br>**vAg**: neg | 3770 |
| | | **Lung (HE)**: several small focal areas of (pv) mononuclear (macrophages, LC, a few deg cells) infiltrates with vasculitis<br>**vAg**: many random, mainly small patches of alveoli with pos AEC | 287929 |
| | | **Brain (HE)**: NHA<br>**vAg**: neg | N/A |
| C4.8 | Omicron, 6 dpi | **Nasal mucosa (HE)**: NHA<br>**vAg**: neg | <LOD |
| | | **Lung (HE)**: several small focal areas of (pv) mononuclear (macrophages, LC, a few deg cells) infiltrates with vasculitis; small focal area with AEC desquamation<br>**vAg**: multiple disseminated, mainly small patches of alveoli with pos AEC | 499774 |
| | | **Brain (HE)**: NHA<br>**vAg**: neg | N/A |
| **Ronapreve treated animals** | | | |
| R1.1 | Delta, 4 dpi | **Nasal mucosa (HE)**: occ degenerate EC<br>**vAg**: some individual and patches of partly deg pos REC and OEC | 73656 |
| | | **Lung (HE)**: mild multifocal activated type II pc, occ syncytial cells and deg AEC; focally IIC; several vessels with vasculitis<br>**vAg**: abundant variably sized, often large disseminated patches of alveoli with pos AEC | 3905724 |
| | | **Brain (HE)**: NHA<br>**vAg**: neg | N/A |
| R1.2 | Delta, 4 dpi | **Nasal mucosa (HE)**: occ degenerate EC<br>**vAg**: a few individual and patches of partly deg pos REC and OEC | 20471 |
| | | **Lung (HE)**: large focal area with activated type II pc, occ syncytial cells, some desquamed AEC and IIC<br>**vAg**: numerous small disseminated patches of alveoli with pos AEC, larger patches in association with focal lesions | 701589 |
| | | **Brain (HE)**: NHA<br>**vAg**: neg | N/A |
| R1.3 | Delta, 4 dpi | **Nasal mucosa (HE)**: occ degenerate EC<br>**vAg**: a few individual and patches of partly deg pos REC and OEC | 75829 |
| | | **Lung (HE)**: one vessel with mild mononuclear infiltration (vasculitis) and pv infitration<br>**vAg**: neg | <LOD |
| | | **Brain (HE)**: NHA<br>**vAg**: neg | N/A |
| R1.4 | Delta, | **Nasal mucosa (HE)**: occ degenerate EC<br>**vAg**: occ individual pos REC and OEC | 197432 |

| | | | |
|---|---|---|---|
| | 4 dpi | **Lung (HE)**: a few small focal areas with activated type II pc, occ syncytial cells and some desquamed AEC<br>**vAg**: numerous small disseminated patches of alveoli with pos AEC, larger patches in association with focal lesions | 2071876 |
| | | **Brain (HE)**: NHA<br>**vAg**: neg | N/A |
| R1.5 | Delta,<br>4 dpi | **Nasal mucosa (HE)**: NHA (small fragment)<br>**vAg**: neg | 42149 |
| | | **Lung (HE)**: a few small focal areas with activated type II pc, occ syncytial cells, some desquamed AEC, occ degenerate cells and some infiltrating LC and NL<br>**vAg**: numerous small disseminated patches of alveoli with pos AEC, larger patches in association with focal lesions | 15340 |
| | | **Brain (HE)**: NHA<br>**vAg**: neg | N/A |
| R1.6 | Delta,<br>4 dpi | **Nasal mucosa (HE)**: NHA (small fragment)<br>**vAg**: neg | <LOD |
| | | **Lung (HE)**: a few small focal areas with activated type II pc, occ syncytial cells and some desquamed AEC<br>**vAg**: several small random patches of alveoli with pos AEC | 342033 |
| | | **Brain (HE)**: NHA<br>**vAg**: neg | N/A |
| R1.7 | Delta,<br>4 dpi | **Nasal mucosa (HE)**: NHA (small fragment)<br>**vAg**: neg | 8373 |
| | | **Lung (HE)**: larger focal area with activated type II pcs, occ syncytial cells, some desquamed AEC and IIC; a few small similar areas<br>**vAg**: numerous small random patches of alveoli with pos AEC, larger patch in association with focal lesion | 28914 |
| | | **Brain (HE)**: NHA<br>**vAg**: neg | N/A |
| R1.8 | Delta,<br>4 dpi | **Nasal mucosa (HE)**: occ deg EC<br>**vAg**: individual and patches of pos, occ deg REC and OEC | 855075 |
| | | **Lung (HE)**: small focal area with activated type II pc, infiltrating macrophages and LC and a few deg cells<br>**vAg**: a few small random patches of alveoli with pos AEC | 300732 |
| | | **Brain (HE)**: NHA<br>**vAg**: neg | N/A |
| R2.1 | Omicron,<br>4 dpi | **Nasal mucosa (HE)**: NHA<br>**vAg**: rare individual intact pos OEC | 14735 |
| | | **Lung (HE)**: mild multifocal IIC<br>**vAg**: neg | <LOD |
| | | **Brain (HE)**: NHA<br>**vAg**: neg | N/A |
| R2.2 | Omicron,<br>4 dpi | **Nasal mucosa (HE)**: NHA<br>**vAg**: neg | 5656 |
| | | **Lung (HE)**: mild multifocal IIC and activated type II pc<br>**vAg**: neg | <LOD |
| | | **Brain (HE)**: NHA<br>**vAg**: neg | N/A |
| R2.3 | Omicron,<br>4 dpi | **Nasal mucosa (HE)**: NHA<br>**vAg**: neg | <LOD |
| | | **Lung (HE)**: mild pv LC dominated mononuclear infiltration<br>**vAg**: very numerous disseminated, mainly small patches of alveoli with pos AEC | 190447 |
| | | **Brain (HE)**: NHA<br>**vAg**: neg | N/A |
| R2.4 | Omicron,<br>4 dpi | **Nasal mucosa (HE)**: NHA<br>**vAg**: neg | 7392 |
| | | **Lung (HE)**: small focal mixed cellular infiltrates<br>**vAg**: very numerous disseminated small to moderately sized patches of alveoli with pos AEC | 614777 |
| | | **Brain (HE)**: NHA<br>**vAg**: neg | N/A |

| | | | |
|---|---|---|---|
| R2.5 | Omicron, 4 dpi | **Nasal mucosa (HE)**: NHA<br>**vAg**: neg | <LOD |
| | | **Lung (HE)**: small disseminated focal LC and macrophage aggregates with a few deg cells (also pv)<br>**vAg**: numerous disseminated, mainly small patches of alveoli of pos alveoli (also around focal aggregates) | 470282 |
| | | **Brain (HE)**: NHA<br>**vAg**: neg | N/A |
| R2.6 | Omicron, 4 dpi | **Nasal mucosa (HE)**: NHA<br>**vAg**: neg | 1287 |
| | | **Lung (HE)**: focal IIC, small areas of patchy vasculitis and pv leukocyte infiltrates with a few deg cells<br>**vAg**: numerous random small patches of alveoli with pos AEC | 1377339 |
| | | **Brain (HE)**: NHA<br>**vAg**: neg | N/A |
| R2.7 | Omicron, 4 dpi | **Nasal mucosa (HE)**: NHA<br>**vAg**: neg | <LOD |
| | | **Lung (HE)**: focal areas of activated type II pneumocytes<br>**vAg**: multiple random small patches of alveoli with pos AEC | 3706590 |
| | | **Brain (HE)**: NHA<br>**vAg**: neg | N/A |
| R2.8 | Omicron, 4 dpi | **Nasal mucosa (HE)**: NHA<br>**vAg**: neg | <LOD |
| | | **Lung (HE)**: NHA<br>**vAg**: neg | 12266 |
| | | **Brain (HE)**: NHA<br>**vAg**: neg | N/A |
| R3.1 | Delta, 6 dpi | **Nasal mucosa (HE)**: occasional deg EC<br>**vAg**: several individual, partly deg pos OEC, some individual pos REC at nasal tip | 5141 |
| | | **Lung (HE)**: multifocal small and delineated dense parenchymal infiltrate, patchy vascular (vasculitis) and pv mononuclear (LC, macrophages) infiltrates<br>**vAg**: multiple disseminated very small patches of alveoli with pos intact AEC; pos cells and pos debris in focal infiltrates | N/A |
| | | **Brain (HE)**: NHA<br>**vAg**: neg | N/A |
| R3.2 | Delta, 6 dpi | **Nasal mucosa (HE)**: NHA<br>**vAg**: rare individual pos OEC | <LOD |
| | | **Lung (HE)**: multifocal small and delineated dense parenchymal infiltrates, patchy vascular (vasculitis) and pv mononuclear (LC, macrophages) infiltrates<br>**vAg**: multiple disseminated very small patches of alveoli with pos intact AEC; pos cells and pos debris in focal infiltrates | 75345 |
| | | **Brain (HE)**: NHA<br>**vAg**: neg | N/A |
| R3.3 | Delta, 6 dpi | **Nasal mucosa (HE)**: NHA<br>**vAg**: rare individual pos OEC | 1738 |
| | | **Lung (HE)**: multifocal small and delineated dense parenchymal infiltrates (some with activated type II pc), occ patchy vascular (vasculitis) and pv mononuclear (LC, macrophages) infiltrates<br>**vAg**: multiple disseminated very small patches of alveoli with pos intact AEC incl. alveoli close to focal infiltrates; pos cells and pos debris in focal infiltrates | 193810 |
| | | **Brain (HE)**: NHA<br>**vAg**: neg | N/A |
| R3.4 | Delta, 6 dpi | **Nasal mucosa (HE)**: NHA<br>**vAg**: a few individual pos REC | 9215 |
| | | **Lung (HE)**: multifocal small and delineated dense parenchymal infiltrates, occ patchy vascular (vasculitis) and pv mononuclear (LC, macrophages) infiltrates<br>**vAg**: a few disseminated very small patches of alveoli with pos intact AEC; pos cells and pos debris in focal infiltrates | 40243 |
| | | **Brain (HE)**: NHA<br>**vAg**: neg | N/A |
| R3.5 | Delta, | **Nasal mucosa (HE)**: NHA<br>**vAg**: neg | <LOD |

| | | | |
|---|---|---|---|
| | 6 dpi | **Lung (HE)**: multifocal small and delineated dense parenchymal infiltrates, occ vasculitis and pv mononuclear (LC, macrophages) infiltrates<br>**vAg**: a few very small patches of alveoli with pos intact AEC; pos cells and pos debris in focal infiltrates | 34217 |
| | | **Brain (HE)**: NHA<br>**vAg**: neg | N/A |
| R3.6 | Delta,<br>6 dpi | **Nasal mucosa (HE)**: NHA<br>**vAg**: neg | <LOD |
| | | **Lung (HE)**: several focal parenchymal and pv mononuclear (LC, macrophages) infiltrates<br>**vAg**: a few very small patches of alveoli with pos intact AEC | 42204 |
| | | **Brain (HE)**: NHA<br>**vAg**: neg | N/A |
| R3.7 | Delta,<br>6 dpi | **Nasal mucosa (HE)**: NHA<br>**vAg**: rare individual pos OEC | 4599 |
| | | **Lung (HE)**: a few small focal parenchymal and pv, macrophage dominated mononuclear infiltrates<br>**vAg**: a few very small patches of alveoli with pos intact AEC; pos cells and pos debris in focal infiltrates | 71974 |
| | | **Brain (HE)**: NHA<br>**vAg**: neg | N/A |
| R3.8 | Delta,<br>6 dpi | **Nasal mucosa (HE)**: NHA, a few deg cells in lumen<br>**vAg**: very rare individual deg pos OEC | 91704 |
| | | **Lung (HE)**: multifocal small and delineated dense parenchymal and occ patchy vascular (vasculitis) and pv mononuclear (LC, macrophages) infiltrates<br>**vAg**: a few very small patches of alveoli with pos intact AEC; pos cells and pos debris in focal infiltrates | 50260 |
| | | **Brain (HE)**: NHA<br>**vAg**: neg | N/A |
| R4.1 | Omicron,<br>6 dpi | **Nasal mucosa (HE)**: NHA<br>**vAg**: neg | <LOD |
| | | **Lung (HE)**: small focal areas with LC, macrophages and activated type II pc, focally with mild patchy vasculitis<br>**vAg**: numerous random small patches of alveoli with mainly intact pos AEC | 2200313 |
| | | **Brain (HE)**: NHA<br>**vAg**: neg | N/A |
| R4.2 | Omicron,<br>6 dpi | **Nasal mucosa (HE)**: NHA<br>**vAg**: neg | 1581 |
| | | **Lung (HE)**: numerous focal, mainly pv areas with LC, macrophages and mild AEC desquamation or activated type II pc, also partly patchy vasculitis and pv leukocyte infiltration<br>**vAg**: numerous disseminated small to moderately sized (areas with AEC desquamation) patches of alveoli with pos AEC | 1430337 |
| | | **Brain (HE)**: NHA<br>**vAg**: neg | N/A |
| R4.3[4] | Omicron,<br>6 dpi | **Nasal mucosa (HE)**: NHA<br>**vAg**: neg | 10104 |
| | | **Lung (HE)**: rare small areas with LC, macrophages and some activated type II pc<br>**vAg**: numerous disseminated small patches of alveoli with pos AEC | 887195 |
| | | **Brain (HE)**: NHA<br>**vAg**: neg | N/A |
| R4.4 | Omicron,<br>6 dpi | **Nasal mucosa (HE)**: NHA<br>**vAg**: neg | 1461 |
| | | **Lung (HE)**: a few small focal areas with LC, macrophages and some activated type II pc, delineated, partly pv and associated with patchy vasculitis<br>**vAg**: numerous disseminated, mainly small patches of alveoli with mainly intact pos AEC | 1118796 |
| | | **Brain (HE)**: NHA<br>**vAg**: neg | N/A |
| R4.5 | | **Nasal mucosa (HE)**: NHA<br>**vAg**: neg | 3804 |

| | | | |
|---|---|---|---|
| | Omicron, 6 dpi | **Lung (HE)**: several slightly delineated small focal areas with LC, macrophages and some activated type II pc, partly pv and associated with patchy vasculitis<br>**vAg**: numerous disseminated, mainly small patches of alveoli with mainly intact pos AEC | 1718363 |
| | | **Brain (HE)**: NHA<br>**vAg**: neg | N/A |
| R4.6 | Omicron, 6 dpi | **Nasal mucosa (HE)**: NHA<br>**vAg**: neg | <LOD |
| | | **Lung (HE)**: several small focal areas with LC, macrophages and some activated type II pc, partly pv and associated with patchy vasculitis<br>**vAg**: numerous disseminated, mainly small patches of alveoli with pos AEC (also in association with focal lesions) | 501190 |
| | | **Brain (HE)**: NHA<br>**vAg**: neg | N/A |
| R4.7 | Omicron, 6 dpi | **Nasal mucosa (HE)**: NHA<br>**vAg**: neg | 3535 |
| | | **Lung (HE)**: some small focal areas with LC, macrophages and some activated type II pc, partly pv and associated with vasculitis, in one area with mild AEC desquamation<br>**vAg**: numerous disseminated, mainly small patches of alveoli with pos AEC | 1703838 |
| | | **Brain (HE)**: NHA<br>**vAg**: neg | N/A |
| R4.8 | Omicron, 6 dpi | **Nasal mucosa (HE)**: NHA<br>**vAg**: neg | <LOD |
| | | **Lung (HE)**: several small focal areas with LC, macrophages and some activated type II pc, partly pv and associated with vasculitis, some areas with mild AEC desquamation<br>**vAg**: several random small patches of alveoli with pos AEC, larger patches in association with focal lesions | 455750 |
| | | **Brain (HE)**: NHA<br>**vAg**: neg | N/A |

**Legend**: AEC – alveolar epithelial cells (type I and II pneumocytes); BEC – bronchiolar epithelial cells; deg – degenerate; EC – epithelial cells; HE – histological features assessed in a hematoxylin-eosin stained section; IIC – increased interstitial cellularity; in – intranasal; LC – lymphocyte; neg – negative; NHA – no histological abnormality; NL – neutrophilic leukocytes; occ – occasional; OEC – olfactory epithelial cells; pa – periarterial; pb – peribronchiolar; pc – pneumocytes; pos – positive; pv – perivascular; REC – respiratory epithelial cells; vAg – viral antigen; N/A – not available; <LOD – below limit of detection.

[1] Day of euthanasia after infection

[2] Copies of viral sgE-RNA/µg of RNA relative to 18S

[3] Remnants of nasal mucosa after sampling of nasal turbinates for PCR.

[4] Animal euthanised at day 3 post infection due to ill health.
